# Synthetic nanobody–SARS-CoV-2 receptor-binding domain structures identify distinct epitopes

**DOI:** 10.1101/2021.01.27.428466

**Authors:** Javeed Ahmad, Jiansheng Jiang, Lisa F. Boyd, Kannan Natarajan, David H. Margulies

**Author notes:** Correspondence to (D.H.M). Equal contributions (Javeed Ahmad, Jiansheng Jiang).

## Abstract

The worldwide spread of severe acute respiratory syndrome coronavirus 2 (SARS-CoV-2) demands unprecedented attention. We report four X-ray crystal structures of three synthetic nanobodies (sybodies) (Sb16, Sb45 and Sb68) bind to the receptor-binding domain (RBD) of SARS-CoV-2: binary complexes of Sb16–RBD and Sb45–RBD; a ternary complex of Sb45–RBD–Sb68; and Sb16 unliganded. Sb16 and Sb45 bind the RBD at the ACE2 interface, positioning their CDR2 and CDR3 loops diametrically. Sb16 reveals a large CDR2 shift when binding the RBD. Sb68 interacts peripherally at the ACE2 interface; steric clashes with glycans explain its mechanism of viral neutralization. Superposing these structures onto trimeric spike (S) protein models indicates these sybodies bind conformations of the mature S protein differently, which may aid therapeutic design.

**One Sentence Summary:** X-ray structures of synthetic nanobodies complexed with the receptor-binding domain of the spike protein of SARS-CoV-2 reveal details of CDR loop interactions in recognition of distinct epitopic sites.

## Main Text

SARS-CoV-2, a β-coronavirus, is remarkable for its high infectivity, rapid worldwide dissemination, and evolution of highly infectious new variants (*1–4*). The virus exploits its trimeric S glycoprotein to adsorb to the host cell-surface receptor, angiotensin converting enzyme (ACE) ACE2 (*5*) resulting in proteolytic processing and conformational changes required for membrane fusion and cell entry (*6*). Understanding the fundamental molecular and cell biology and chemistry of the viral life cycle and the nature of the host immune response, offers rational avenues for developing diagnostics, therapeutics, and vaccines (*7, 8*). Exploring the detailed structures of anti-viral antibodies can provide critical understanding of the means to attenuate viral adsorption and entry, preventing or retarding ongoing infection and communal spread. An evolving database of X-ray and cryo-EM structures of the SARS-CoV-2 S and its interactions with ACE2 or various antibodies contributes to the design of effective antibodies or immunogens (*9*). Recent studies indicate the value of single domain antibodies derived from camelids (nanobodies) (*10*) or camelid-inspired synthetic libraries (sybodies) (*11*), and the value of generating multivalent constructs (*12*) for effective treatment (*11*). Many properties of nanobodies make them well suited for structural studies and drug development (*13*).

Here, we take advantage of available sequences of three SARS-CoV-2 RBD-directed sybodies – Sb16, Sb45, and Sb68 (previously designated Sb#16, Sb#45, and Sb#68 (*14*)). We describe binding studies and X-ray structures of complexes of these with the RBD, and also the structure of Sb16 unliganded. The sybodies had been shown to be effective inhibitors of the ACE2–RBD interaction (*14*), and neutralizers of viral infectivity (*14*). These sybodies (see Supplementary Materials and Methods) behaved as monomers by size exclusion chromatography (SEC) (*15*) (Figure S1), and we confirmed their activity in binding to the re-engineered RBD and S using surface plasmon resonance (SPR) (Figure S2). All three sybodies bind to surface immobilized RBD with *K*_D_ values of 0.038 to 0.77 μM (Figure S2A to S2D) – measurements that are similar to those determined using RBD-YFP or RBD-Fc molecules by related techniques (*14*). Binding of Sb16, Sb45, and Sb68 to S consistently revealed lower affinities, in the range of 0.07 to 2.6 μM (Figure S2E to S2H). Experiments using SEC of premixed solutions of sybodies and RBD confirmed that all three sybodies bound the RBD (Figure S3). RBD consistently eluted at 13 min. Mixtures of RBD with Sb16 or Sb45 eluted at ~11.3 min and with Sb68 at ~11.7 min consistent with complex formation. For unliganded Sb16, the large change in elution time suggests that its RBD binding site is that which interacts with the chromatographic column matrix (Figure S3).

To gain insight into the precise topology of the interaction of each of the three sybodies with the RBD, we determined crystal structures of these complexes. We obtained crystals of several complexes: Sb16–RBD, Sb45–RBD, and the ternary Sb45–RBD–Sb68; and of Sb16 alone. These crystals diffracted X-rays to resolutions from 2.1 to 2.6 Å (Table S1). After molecular replacement, model building, and crystallographic refinement (see Materials and Methods), we obtained structural models with *R*_work_/*R*_free_ (%) of 25.4/28.4, 18.6/21.6, 20.6/25.5 and 22.5/25.6, respectively, that satisfied standard criteria for fitting and geometry (Table S1). Illustrations of the quality of the final models as compared with the electron density maps are shown in Figure S4.

The structure of the RBD domain of these complexes (Figure 1A and 1B) revealed little difference between insect-expressed (*16*) and bacteria-expressed and refolded RBD. Each of the sybodies has a barrel of two β-sheets stabilized by a single disulfide-linked loop of 75 or 76 amino acids characteristic of an IgV fold (*17, 18*). The Sb16–RBD complex (Figure 1A and 2A) illustrates that CDR2 (residues 50-60) and CDR3 (residues 98-106) bestride the saddle-like region of the ACE2-binding surface of the RBD (see sequence alignment in Figure 1E). Sb16 angulates over the RBD by 83°. However, Sb45 (Figure 1B and 2B) straddles the RBD saddle in the opposite orientation, at an angle of −36°, and frames the interface with CDR2 (residues 50-59) and CDR3 (residues 97-111). CDR1s of both sybodies (residues 27-35) lie between the CDR2 and CDR3 loops. Superposition of the two structures, based on the RBD, emphasizes the diametrically opposite orientation of the two (Figure 1C), revealing that the CDR2 of Sb16 and CDR3 of Sb45 recognize the same epitopic regions.

**Fig. 1.**
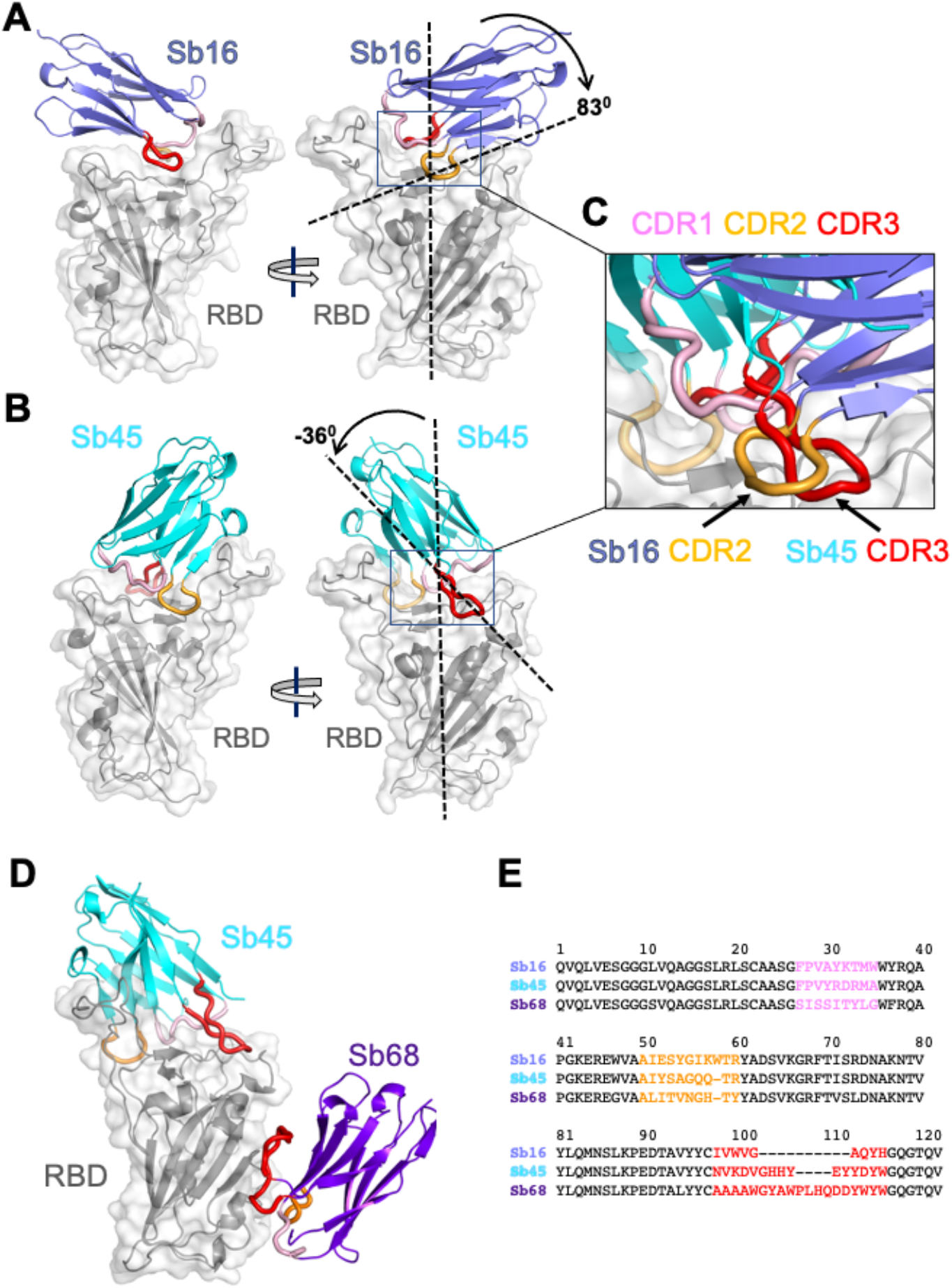
Overall structures of Sb16, Sb45 and Sb68 complexes with SARS-CoV-2 RBD. Ribbons (sybodies) and ribbons plus surface (RBD) representations of the complex of (A) Sb16 (slate) with RBD (grey) (7KGK); (B) Sb45 (cyan) with RBD (7KGJ), and (C) Sb45 and Sb68 (purple) with RBD (7KLW). Sb16-RBD and Sb45-RBD, superimposed based on the RBD are shown in (D) to highlight CDR loops, which are color coded as indicated. The CDR2 of Sb16 and CDR3 of Sb45 interact similarly with the RBD surface.

**Fig. 2.**
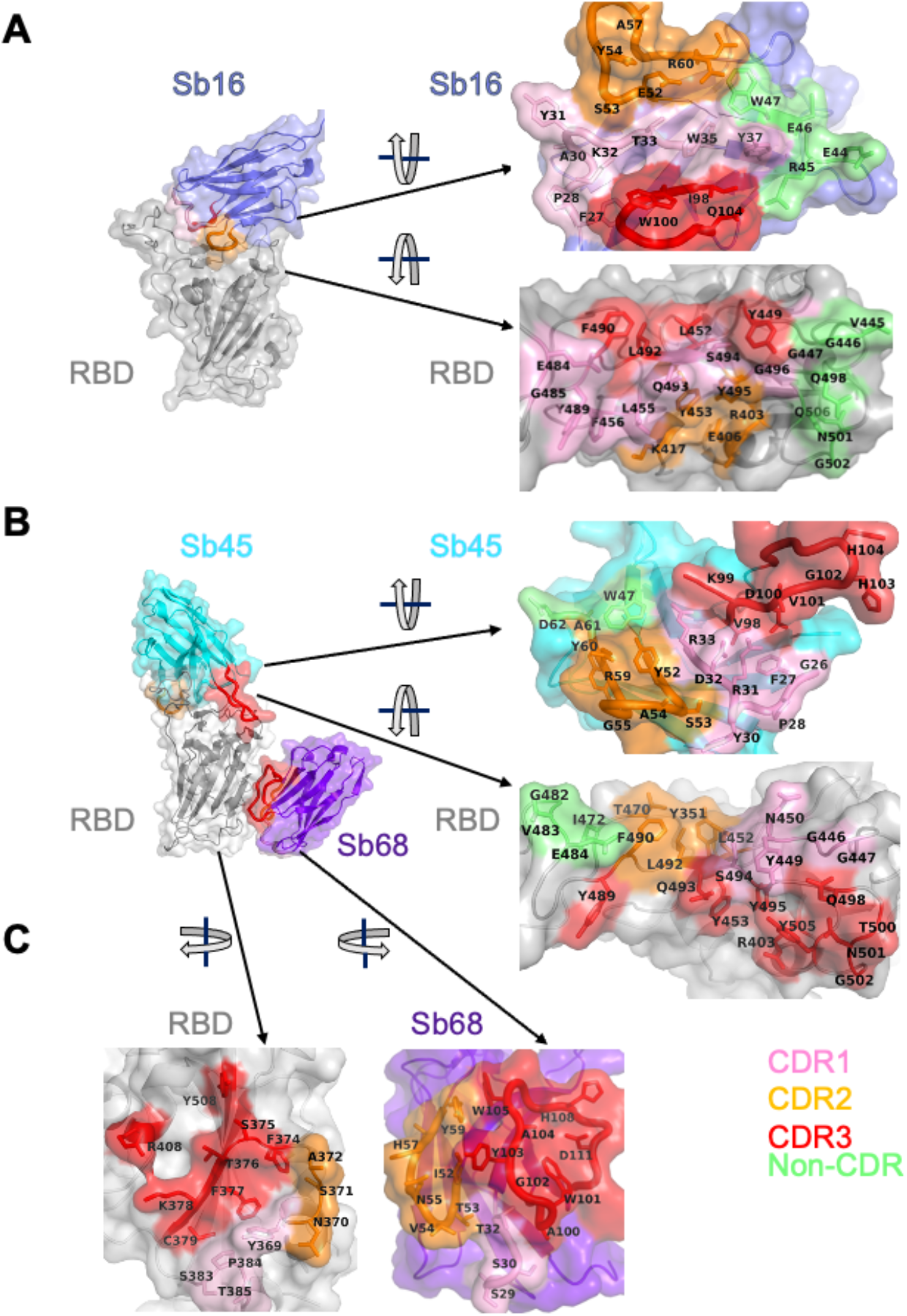
Interface and interaction of (A) Sb16-RBD, (B) Sb45-RBD and (C) Sb68-RBD. (Individual contacting residues are listed in TableS2 in Supplemental Materials). CDR1, CDR2, CDR3 regions are painted pink, orange and red respectively. Additional non-CDR region contacting residues are colored lime. On the RBD surface, the epitopic residues that contact the sybodies are colored according to the sybody CDR.

Exploring conditions using mixtures of two or three sybodies and the RBD, we obtained crystals and solved the structure of a ternary complex consisting of Sb45–RBD–Sb68 at 2.6 Å (Table S1 and Figure 1D). The refined model revealed Sb45 and Sb68 interacting at two different faces of the RBD (Figure 1D and 2B). Here, Sb45 binds in an identical orientation to that observed in the binary Sb45–RBD structure (RMSD of superposition, 0.491 Å for 1981 atoms), but Sb68 addresses a completely different face of the RBD – similar to that bound by Fab of CR3022 on RBD of SARS-CoV-2 (*19*) and by V_HH_72 on RBD of SARS-CoV-1 (*20*). Of particular interest, whereas Sb45 CDR2 and CDR3 span the RBD saddle as noted above, the distinct contacts of Sb68 to the RBD are through the longer CDR3, with only minor contributions from CDR1 and CDR2. Walter et al visualized similar distinct interactions in cryo-EM maps of two sybodies (Sb15 and Sb68) bound to S protein with local resolution of 6-7 Å (*14*).

Scrutiny of the different interfaces provides insights into the distinct ways each sybody exploits its unique CDR residues for interaction with epitopic residues of the RBD (Figure 2). Both Sb16 and Sb45 use longer CDR2 and CDR3 to straddle the RBD, positioning CDR1 residues over the central crest of the saddle (Figure 2A and 2B). Also, several non-CDR residues (Y37, E44, R45, E46, and W47 for Sb16; and W47 for Sb45), derived from framework 2 (*21*), provide additional contacts to the RBD. The interface of Sb68 with RBD (Figure 2C) is quite different, predominantly exploiting eight CDR3, six CDR2, and four CDR1 residues, along with non-CDR residues at the interface. (Table S2 lists all individual contacts between each sybody and the RBD).

To evaluate the structural basis for the ability of these three sybodies to block the interaction of RBD with ACE2, we superposed each of the three sybody–RBD structures onto the ACE2–RBD structure and examined the steric clashes (Figure 3A). Sb16 and Sb45 directly impinge on the ACE2 binding site, offering a structural rationale for their viral neutralization capacity (*14*). Sb68, which also blocks viral infectivity, binds to RBD at a site which appears to be noncompetitive for ACE2 binding. The carbohydrate at ACE2 residues N322 and N546 provides an explanation (Figure 3A, and below).

**Fig. 3.**
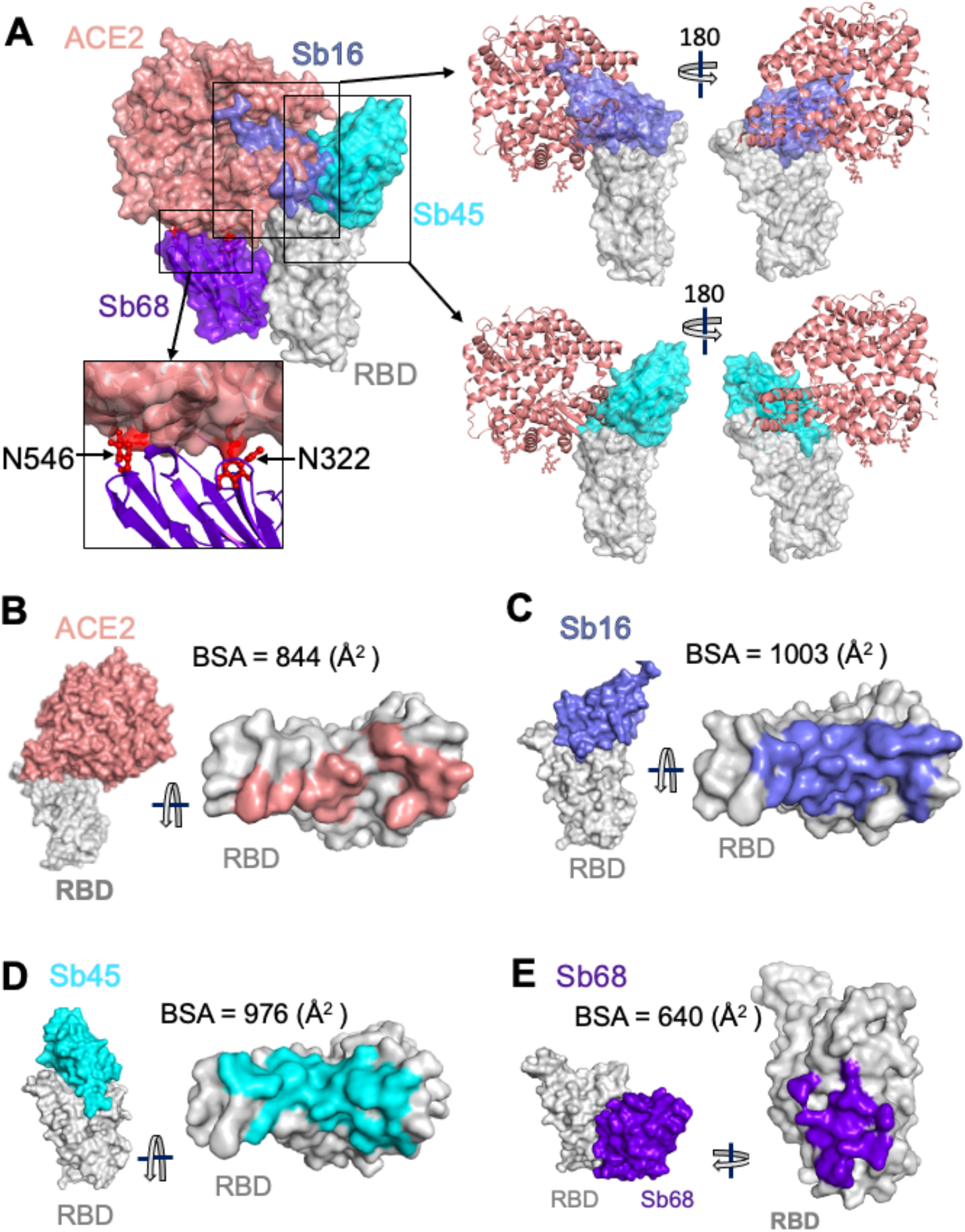
Sybodies compete with ACE2 for RBD binding. (A) Sb16 (slate), Sb45 (cyan) and Sb68 (purple) – RBD complexes were superposed on the ACE2—RBD structure (salmon) (6M0J) based on the RBD. Sb16 is buried inside ACE2; Sb45 is partially buried in ACE2; and Sb68 has major clashes with two N-glycan sites (N322 and N546) of ACE2. (B) Epitopic areas (on RBD) captured by ACE2 (salmon), BSA = 844 (Å^2^); (C) by Sb16 (slate), BSA = 1003 (Å^2^); (D) by Sb45 (cyan), BSA = 976 (Å^2^); and (E) by Sb68 (purple), BSA = 640 (Å^2^).

To compare the epitopic areas captured by these sybodies, we evaluated the buried surface area (BSA) interfaces between RBD and ACE2 or the sybodies. The BSA at the ACE2– RBD, Sb16–RBD, Sb45–RBD, and Sb68–RBD interfaces are 844 Å^2^, 1,003 Å^2^, 976 Å^2^, and 640 Å^2^, respectively (Figure 3B to 3E). Sb16 and Sb45 capture more surface area than ACE2 or other published nanobody or sybody–RBD complexes (see Table S3). The interface with Sb68 is the smallest (640 Å^2^) (Figure 3E). The total BSA captured by Sb45 and Sb68 in the ternary complex is 1,650 (1,010 plus 640) Å^2^ (Table S4) and is consistent with the view that a linked bispecific sybody, as described by Walter et al (*14*), would exert strong avidity effects. Table S3 summarizes these BSA values and those of other nanobody–RBD interactions.

A reasonable explanation for the ability of Sb68 to block the ACE2–RBD interaction arises on inspection of the sites where Sb68, bound to the RBD, might clash with ACE2. Scrutiny of a superposition of Sb68–RBD with ACE2–RBD reveals several areas of steric interference. Sb68 loop 40-44 clashes with amino acid side chains of ACE2 (residues 318-320 and 548-552), loop 61-64 with ACE2 N322 carbohydrate, and loop 87-89 (a 3,10 helix) with ACE2 N546 carbohydrate as well as residues 313 and 316-218 (Figure 3A). The ACE2 used in the crystallographic visualization of ACE2–RBD (*22*) was expressed in *Trichoplusia ni* insect cells, which produce biantennary N-glycans terminating with N-acetylglucosamine residues (*23, 24*). Electron density was observed only for the proximal N-glycans at residues N322 and N546, but larger, complex, non-sialylated, biantennary carbohydrates have been detected in glycoproteomic analysis of ACE2 in mammalian cells (*25*). These are highly flexible carbohydrates adding greater than 1500 Da at each position, so are larger than the single carbohydrate residues visualized in the crystal structure. Additionally, molecular dynamics simulations of RBD–ACE2 implicated the direct interaction of carbohydrate with the RBD (*26*). Thus, the ability of Sb68 to impinge on ACE2 interaction with RBD likely involves the steric clash of the N322- and N546-linked glycans.

We also obtained a 2.1 Å structure of free Sb16 (Figure S5). Remarkably, the CDR2 of Sb16 shows Y54 in starkly different positions in the unliganded structure as compared to the complex: the Cα carbon is displaced by 6.0 Å, while the Oη oxygen of Y54 is 15.2 Å distant, indicative of dynamic flexibility.

To gain additional insight into the structural consequences of the interactions of each of these sybodies with a trimeric S protein, we superposed each of the individual sybody–RBD complexes on each of several cryo-EM-determined models of S, including examples of different combinations of RBD orientation: three-down (6XEY (*27*)), one-up, two-down (6Z43 (*28*)), two-up, one-down (7A29 (*29*)), and three-up (7JVC (*30*)) (see Figure 4). Both Sb16 and Sb45 may dock on each of the three RBDs in the trimeric S in any of the four configurations, without any apparent clash (Figure 4A, 4B). However, Sb68 could not be superposed without clashes to any RBD of the three-down or to the one-up two down position. The only permissible superpositions were to two in the two-up, one-down (Figure 4C); and to all three in the three-up position (Figure 4C). For paired sybodies, either Sb16 and Sb68 or Sb45 and Sb68, superposition was possible without clashes, with two or more RBDs in the up conformation (Figure 4D and 4E). Walter et al (*14*) suggested that a covalent bispecific Sb15–Sb68 reagent could bind S in both the two-up and three-up configurations, based on cryo-EM maps of complexes of S with Sb15 and Sb68, with local resolution in the range of 6-7 Å. It appears that Sb16 binds to S in an orientation similar to, but in detail distinct from that of Sb15. This analysis demonstrates an advantage of the small size of sybodies or nanobodies in accessing epitopic regions of S.

**Fig. 4.**
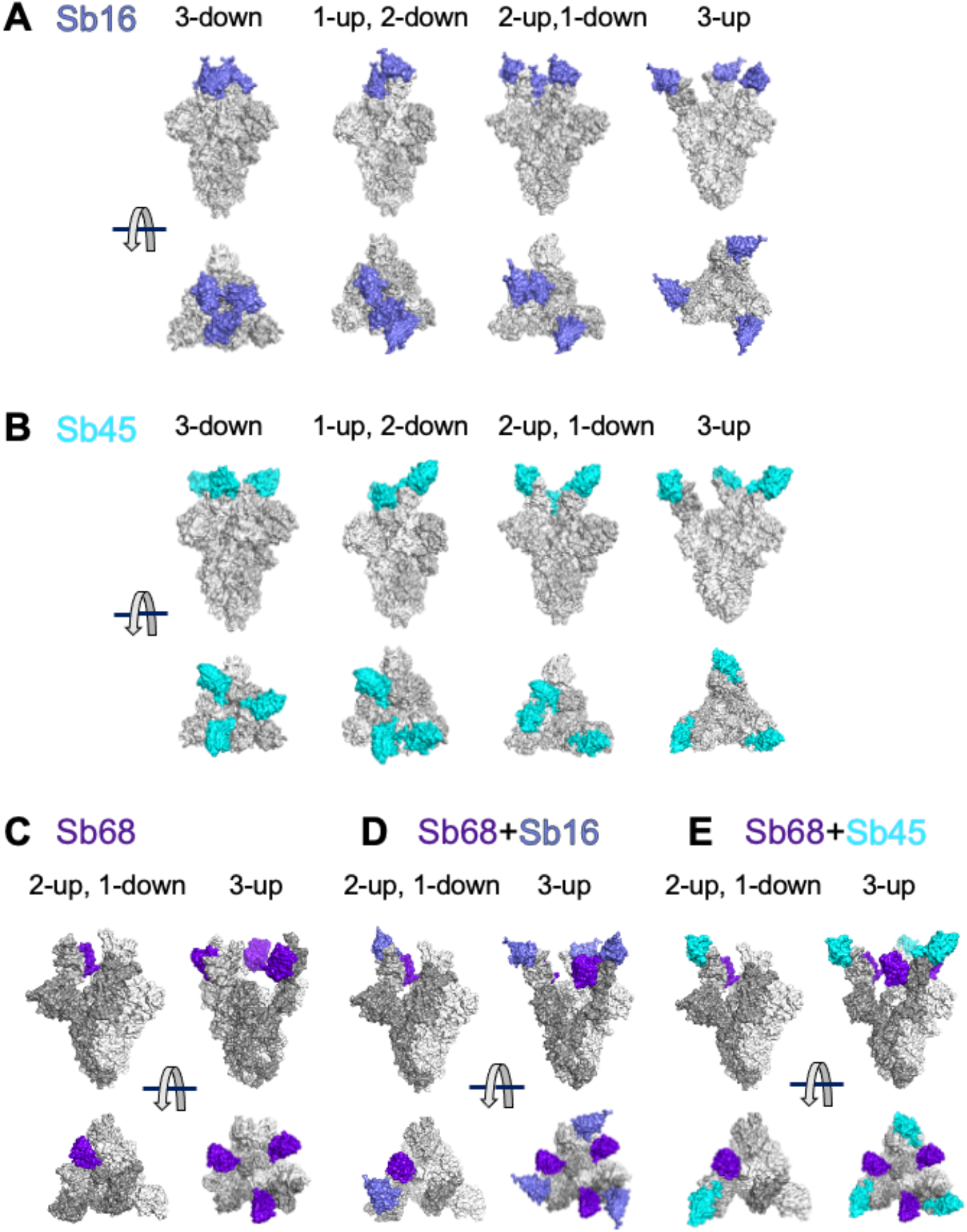
Superposition of complexes on spike models reveals accessibility of sybodies. Superposition of (A) Sb16-RBD (slate) on spike (6XEY: 3-down; 6Z43: 1-up and 2-down; 7A29: 2-up and 1-down; 7JVC: 3-up); (B) Sb45-RBD (cyan); and (C) Sb68-RBD (purple), there is no accessible surface for Sb68 on 3-down of spike; (D) Sb68 and Sb16 on RBD; (E) SB68 and Sb45 on RBD.

Barnes et al (*31*) categorized a host of anti-S and anti-RBD Fabs into four classes (1-4) based on the location of the footprint, and whether the Fab has access to either the up only or up and down configuration of the RBD in the context of the full trimer (Figure S6A). Xiang et al (*12*) categorized anti-RBD nanobodies into five epitopic regions (I-V) (Figure S6A). By superposition, Sb16 would clash with the light chain of the B38 Fab (7BZ5), as in class 1, but it also clashes with the heavy chain of COVA2-39 (7JMP), as in class 2 (Figure S6B). Sb45 clashes effectively with the heavy chain of COVA2-39, and thus appears to be closer to a “true” class 2 sybody (Figure S6C). Both Sb16 and Sb45 are capable of binding the RBD of S in either the up or down position, a defining characteristic of class 2. By contrast, Sb68 competes mostly with the CR3022 heavy chain (6W41), V_HH_72 (6WAQ) (*20*) and V_HH_-U (7KN5) (*32*) placing it in class 4. Overall, our structural studies not only define the Sb16, Sb45, and Sb68 epitopes at high resolution, they suggest that a battery of sybodies or nanobodies have the potential to saturate the available RBD surface.

The significance of the ternary structure of Sb45–RBD–Sb68 (7KLW) is confirmed in a recent paper (*32*). Koenig et al determined a ternary nanobody structure of V_HH_-E–RBD–V_HH_-U (7KN5) which illustrates the binding to two distinct epitopic sites. Superposition of Sb45–RBD– Sb68 on V_HH_-E–RBD–V_HH_-U indicates that Sb45 and V_HH_-E represent class 2 in recognizing the epitope region but do so in different orientations (Figure S7A, middle panel). Sb45 uses both long CDR2 and CDR3 loops riding along both sides of RBD surface, while V_HH_-E uses a long CDR3 loop engaging one side of RBD surface.

Recently, several SARS-CoV-2 spike variants have been isolated and characterized with respect to their infectivity and severity of disease. The UK-SARS-CoV-2 variant has multiple substitutions including N501Y in the RBD (*1*). This is expected to impinge on the peripheral aspect of the footprint of Sb16 and Sb45 but would have no effect on the Sb68 site. Thus, precise mapping of anti-RBD antibody, nanobody, and sybody epitopes, especially for those that are developed for clinical trials, has implications not only for mechanistic understanding of the interactions of the RBD with ACE2, but also for evaluating the potential susceptibility of newly arising viral variants to currently administered vaccines and antibodies.

## Acknowledgments

We appreciate the help of Joy (Huaying) Zhao and Peter Schuck, NIBIB, NIH in analyzing SPR data, and thank Peter Sun, NIAID, NIH for access to his program, HINGE. We thank Barney Graham, NIAID, NIH for plasmids used in initial aspects of the work, and Apostolos Gittis, NIAID, NIH, for help in protein characterization. We appreciate the advice of Michael Mage and D. K. Taylor during this work.

## Author contributions

J.A., J.J., K.N., and D.H.M. conceived the project; J.A., K.N., and L.F.B.engineered constructs, purified protein, and performed binding analyses; J.J. screened for crystals, processed X-ray data, and refined the structures; all authors contributed to the final manuscript.

## Data and materials availability

All data are included in the paper or in the supplementary material. X-ray structure factors and coordinates are deposited at the protein data bank (www.pdb.org) under accession numbers 7KGK, 7KGJ, 7KLW and 7KGL for Sb16–RBD, Sb45–RBD, Sb45–RBD–Sb68, and Sb16 respectively.

## Supplementary Materials

## Materials and Methods

### Subcloning, expression and purification of RBD, spike, and sybody proteins

The sequences encoding the RBD of the SARS-CoV-2 spike protein (amino acids 333 to 529) were subcloned into pET21b(+), (Novagen) via *Nde*I and *Eco*RI restriction sites, using pcDNA3-SARS-CoV-2-RBD-8his (Addgene #145145, (*33*)) as template. The primers used were forward primer, 5’-TGCAGTCATATGAATCTTTGTCCGTTCGGTGAG and reverse primer, 5’-TGCAGTGAATTCTCACCCTTTTTGGGCCCACAAACT. The RBD was expressed as inclusion bodies in *E. coli* strain BL21(DE3) (Novagen). Expression, isolation of inclusion bodies, denaturation and reduction was done in 6 M guanidine hydrochloride and 0.1 mM DTT as described elsewhere (*34*). Briefly, refolding was carried out in a refolding buffer supplemented with oxidized and reduced glutathione and arginine for 3 days at 4 °C followed by dialysis against HEPES buffer (25 mM HEPES, pH 7.3, 150 mM NaCl). Concentrated and filtered protein was analyzed by size-exclusion chromatography on a Superdex 200 10/300 GL column (GE Healthcare) equilibrated with HEPES buffer. The peak corresponding to 24 kDa (monomeric) protein was collected, concentrated and further purified by ion-exchange chromatography on Mono-Q^®^ (Cytiva).

Plasmids pSb-init encoding sybodies Sb16, Sb45. and Sb68 (Addgene #153524, #153526, and #153527, respectively) were originally reported by Walter et al (*35*) and generously made available. All plasmids were verified by DNA sequencing. Purification of the recombinant proteins from the periplasm of *E. coli* MC1061 was based on a protocol described elsewhere (*36*). Briefly, *E. coli* MC1061, transformed with a sybody-encoding plasmid, was grown in Terrific Broth (TB) medium (Gibco) supplemented with 25 μg/ml chloramphenicol, at 37 °C with shaking at 160 rpm for 2 hrs. The temperature was then decreased to 22 °C until A_600_ reached 0.5. Protein expression was induced by addition of L-(+)-arabinose (Sigma) to a final concentration of 0.02% (w/v) and expression continued overnight at 22 °C and 160 rpm. The next day cells were collected by centrifugation at 2000 x *g* for 15 minutes. The cell pellet was then washed twice in PBS and resuspended in periplasmic extraction buffer (50 mM Tris/HCl pH 8.0, 0.5 mM EDTA, 0.5 μg/ml lysozyme, 20% w/v sucrose (Sigma)) at 4 °C for 30 min followed by addition of TBS (pH 8.0) and 1 mM MgCl_2_. Cells were then centrifuged at 10,000 rpm (Fiberlite™ F21-8 x 507 Fixed Angle Rotor) for 30 min. Following transfer of the supernatant to a fresh tube, imidazole was added to a final concentration of 10 mM. Ni-NTA resin (Qiagen) equilibrated with TBS was added to the supernatant and incubated for 1 hr at RT with mild agitation. The resin was collected, washed three times with buffer supplemented with 30 mM imidazole and sybody proteins were eluted with 300 mM imidazole in TBS.

Plasmid encoding spike HexaPro (designated “S” throughout) was procured from Addgene (#154754) and transfected into Expi293F™ cells (ThermoFisher Scientific) using manufacturer’s protocol. Briefly, Expi293F™ cells were seeded to a final density of 2.5-3 × 10^6^ viable cells/ml and grown overnight at 37 °C in Expi293™ Expression Medium (Gibco). The following day, cell viability was determined, and cell density was adjusted to 3 × 10^6^ viable cells/ml with fresh, prewarmed Expi293™ Expression Medium. Transfection was then done as per manufacturer’s instructions using 1 μg/ml plasmid DNA. Cultures were grown for 6 days following transfection and supernatant was collected, filtered through a 0.22 μm filter and passed over Ni-NTA resin for affinity purification. Further purification was accomplished by size-exclusion chromatography using a Superose 6 10/300 GL column (Cytiva) in a buffer consisting of 2 mM Tris pH 8.0, 200 mM NaCl.

### Preparative and analytical size-exclusion chromatography

Sybodies purified by Ni-NTA affinity chromatography were concentrated using Amicon 10K MWCO concentrators and purified on a Sepax SRT-10C SEC100 column at a flow rate of 1 ml/min. Monomeric sybodies elute at a retention volume of 11–12.5 ml from the Sepax SRT-10C SEC100 column. Monomeric peak fractions were collected and analyzed by SDS-PAGE. Analytical SEC of RBD sybody complexes was performed on a Shodex KW-802.5 column at a flow rate of 0.75 ml/min in TBS buffer (pH. 8.0). (The interaction of individual sybodies with the column matrix is a well-documented phenomenon (*36*)).

### Surface Plasmon Resonance

SPR experiments were performed on a Biacore T200 (Cytiva) at 25 °C in 10 mM HEPES pH 7.2, 150 mM NaCl, 3 mM EDTA, 0.05% Tween-20. RBD was immobilized on a series S CM5 sensor chip (Cytiva) by amine (NHS/EDC) coupling to flow cells. For background subtraction a reference cell was mock coupled. Binding and kinetic studies were performed multiple times for each sybody. Graded and increasing concentrations of SB16, SB45 and SB68 were injected over the RBD-immobilized surface at a flow rate of 30 μl/min with an association time of 120 s and dissociation time of 2000 s. Binding data were analyzed by surface site affinity distribution analysis by EVILFIT (*37, 38*). In general, these values were consistent with fits to the Langmuir binding equation for a 1:1 interaction model using Biacore T200 Evaluation Software v3.0, but revealed better statistics.

### Thermal stability

Thermal melt analysis of the recombinant proteins was performed in triplicate in 96-well plates in a QuantStudio 7 Flex real time PCR machine (Applied Biosystems). Each well contained 2-4 μg protein in buffer (25mM TRIS pH 8, 150 mM NaCl) and 5x Sypro Orange (Invitrogen, stock 5000x) in a total volume of 20 μl. Following an initial two-minute hold at 25 °C, the plate was heated to 99 °C at a rate of 0.05 °C/sec. Data were analyzed with Protein Thermal Shift Software v1.3 (Invitrogen) to obtain T_m_ values for RBD, S, Sb16, Sb45, and Sb68 (Figure S8).

### Crystallization, data collection, structure determination and crystallographic refinement

Purified sybodies (Sb16, Sb45 and Sb68) and RBD were mixed in approximate 1:1 molar ratio to a final concentration of 8 mg/ml. The protein mixtures were incubated on ice for 1 hour prior to screening. Initial screening for crystals was carried out using the hanging drop vapor diffusion method using the Mosquito robotic system (sptlabtech.com). Crystals of SB16-RBD and SB45-RBD complexes and Sb16 alone were observed within one week using Protein Complex (Qiagen) and Wizard Classic 4 (Rigaku). Conditions for Sb16–RBD were either 0.1M HEPES pH 7.0, 15% PEG 20000, or 0.1M HEPES pH 7.0, 18% PEG 12000; and for Sb45–RBD was 18% PEG 12000 and 12% PEG 8000, 0.1 M HEPES pH 7.5, 0.2 M NaCl. Sb16 alone crystallized in 20% PEG 4000, 0.1 M MES pH 6.0, 0.2 M LiSO4. We also screened mixtures of two or three sybodies with RBD. Crystals of Sb45–RBD–Sb68 were obtained after one month following mixing the three proteins in an equimolar ratio in 10% PEG 8000, 0.1M sodium cacodylate pH 6.0.

Crystals of protein complexes were optimized with slight adjustments of the concentration of PEG components. Crystals were cryoprotected in mother liquor containing 5% ethylene glycol and 5% glycerol and flash frozen in liquid nitrogen for data collection. Diffraction data were collected at the Southeast Regional Collaborative Access Team (SER-CAT) beamline 22ID-D at the Advanced Photon Source, Argonne National Laboratory and data were processed with XDS (*39*). Multiple data sets were collected for the protein complexes from 2.3-3.2 Å resolution. The initial model of Sb16 and Sb45 for the molecular replacement search were built by the MMM server (manaslu.fiserlab.org/MMM (*40*)), using the heavy chain V domain and the RBD of the Fab B38–RBD complex (PDBid: 7BZ5) (*41*). The initial model of Sb68 for molecular replacement was built based on the V_H_ domain of 7BZ5. Molecular replacement solutions were found using Phaser (*42, 43*). Subsequent refinements were carried out with PHENIX (*44*). CDR loops were manually rebuilt by fitting to the electron density maps with Coot (*45*). In particular, Sb68 CDR loops were deleted before refinement and built in manually based on electron density maps. Illustrations and calculations of superpositioned models were prepared in PyMOL (The PyMOL Molecular Graphics System, Version 2.4.0 Schrödinger, LLC). Calculation of hinge relationships of domains was accomplished with HINGE (https://niaidsis.niaid.nih.gov/hinge.html), provided courtesy of Peter Sun, NIAID. Buried surface area (BSA) calculations were performed with PISA (https://www.ebi.ac.uk/pdbe/pisa/).

The final structures for the RBD-SB16 and RBD-SB45 complexes showed *R*_work_/*R*_free_ (%) 25.4/27.7 and 18.6/21.6 respectively, and for SB16 alone with *R*_work_/*R*_free_ 22.4/25.9. Data collection and structure refinement statistics are provided in Table S1.

**Fig. S1.**
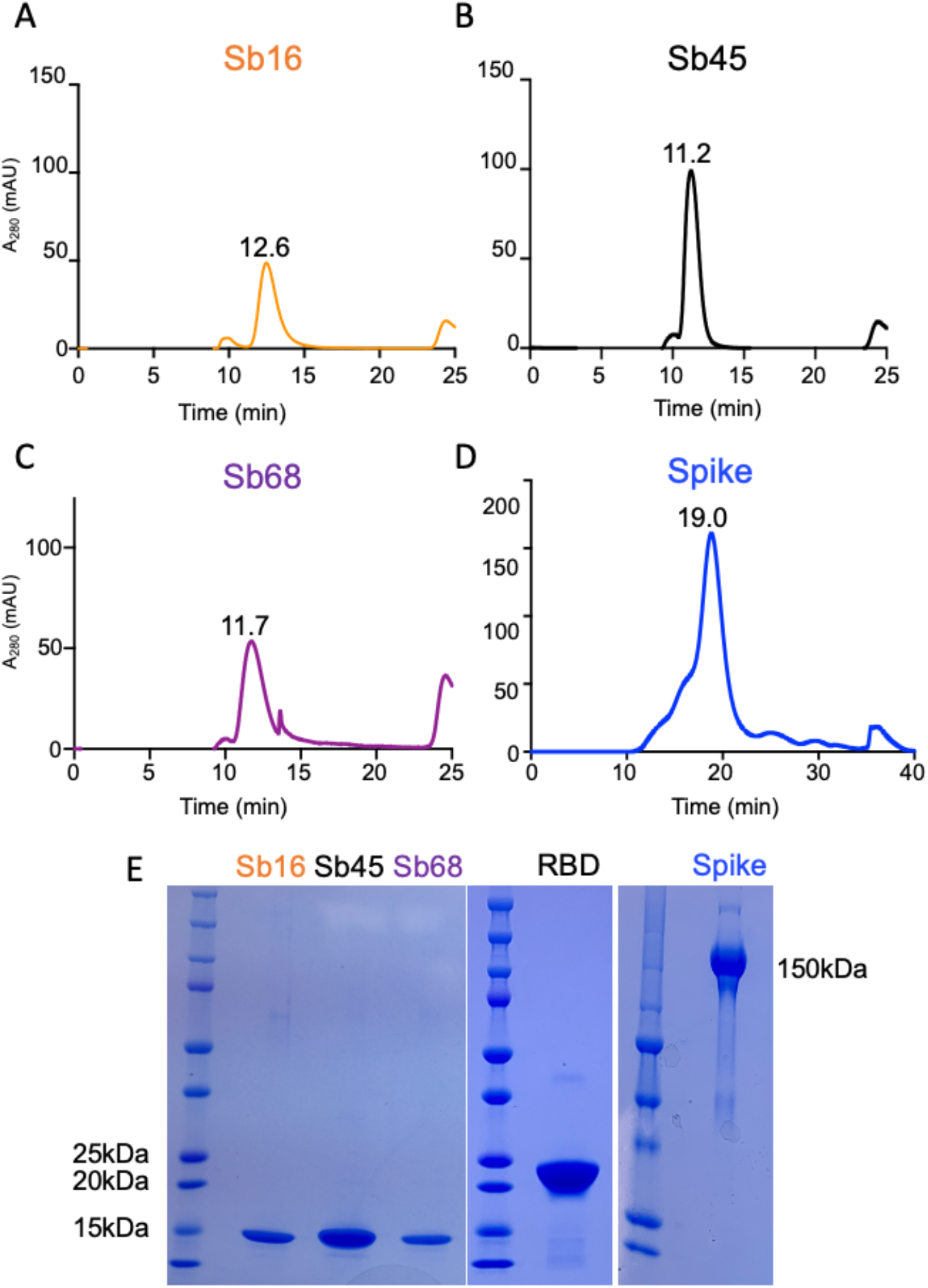
SEC profiles and purification of sybodies, RBD, and spike. (A, B, C) Monomeric sybodies, as indicated, were purified on SRT-10C-SEC100 columns. Elution time of each sybody is indicated above each peak. The *y* axis represents A_280_ nm absorbance units (mAu). (D) SEC profile of trimeric spike protein (Superose™ 6 10/300 GL. (E) SDS-PAGE image of purified sybodies, RBD and spike protein.

**Fig. S2.**
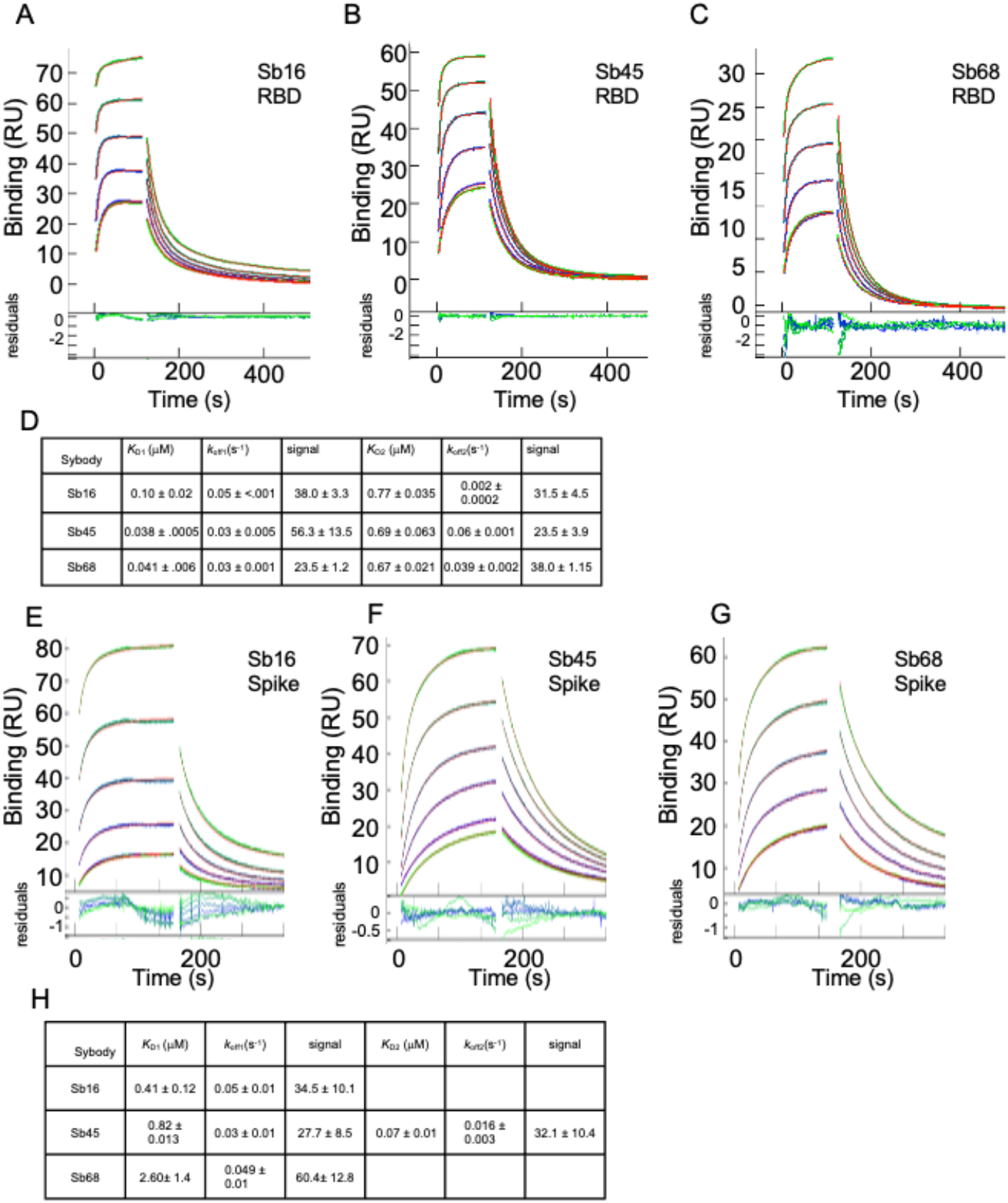
Sybodies bind RBD with *K_D_* values in the nanomolar range. RBD (A, B, C) or S protein (E, F, G) was coupled to a biosensor chip as described in Materials and Methods. Graded concentrations (31 to 500 nM) were flowed over the coupled surfaces (from t=0 to t= 120 s, followed by buffer washout) and net RU signal (compared to mock-coupled surface) was measured by SPR for Sb16 (A, E); Sb45 (B, F); and Sb68 (C, G). Summary of triplicate mean ± SD determinations is shown in Tables (D, H). Global analysis (curve fits in red, residuals below the curves) was accomplished using EVILFIT (*37, 38*), and the major components of binding are shown.

**Fig. S3.**
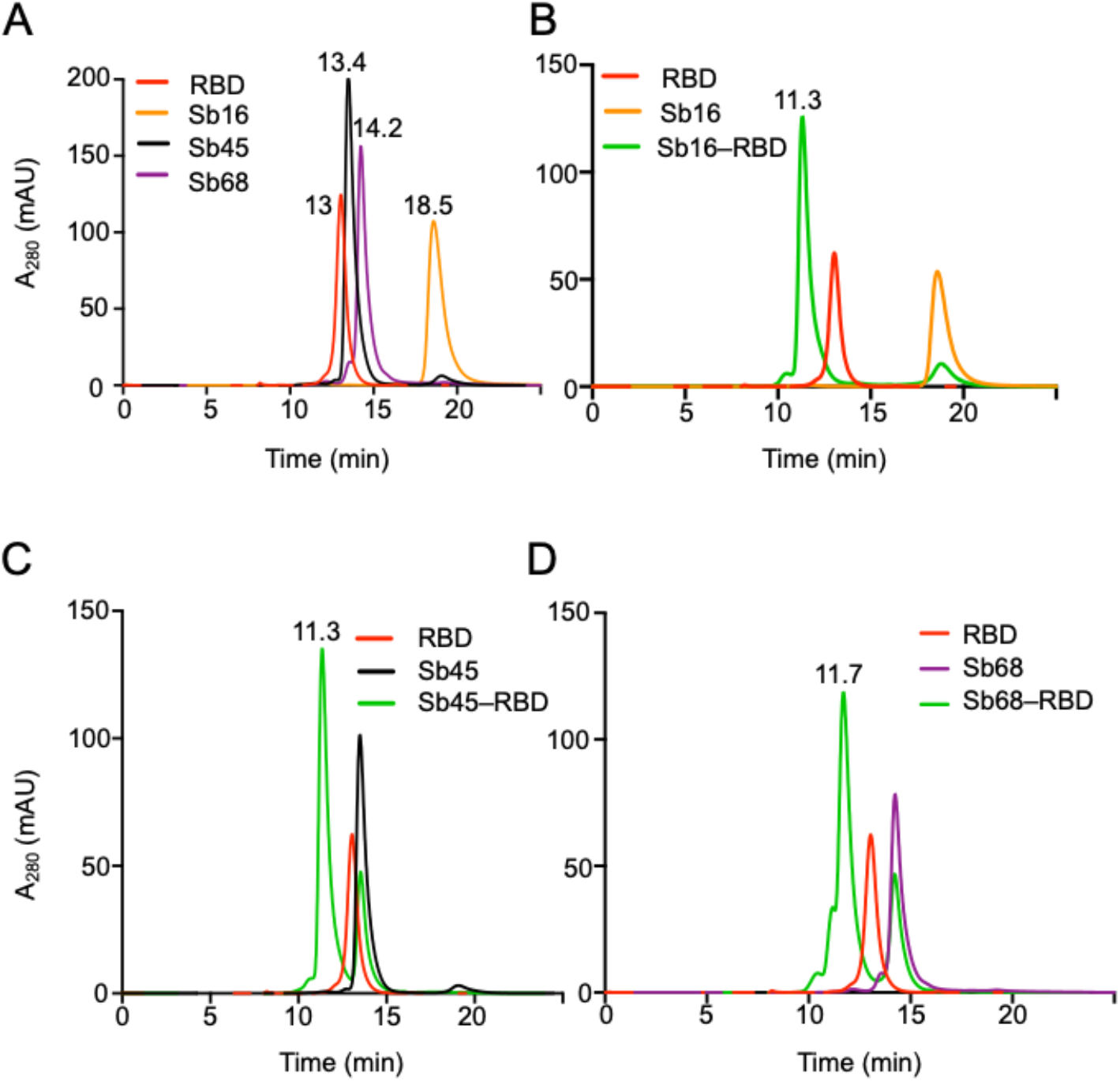
SEC profiles reveal direct interaction of sybodies with RBD. (A) Sybodies (50 μg) and RBD (50 μg) were injected onto a Shodex-KW-802.5 column (0.75 ml/min) in 100 μl individually and elution times are shown. Sb16 and RBD (B), Sb45 and RBD (C), or Sb68 and RBD (D) were mixed in equal concentrations (50 μg in 100 μl), incubated at 4 °C overnight and then analyzed on the Shodex column.

**Fig. S4.**
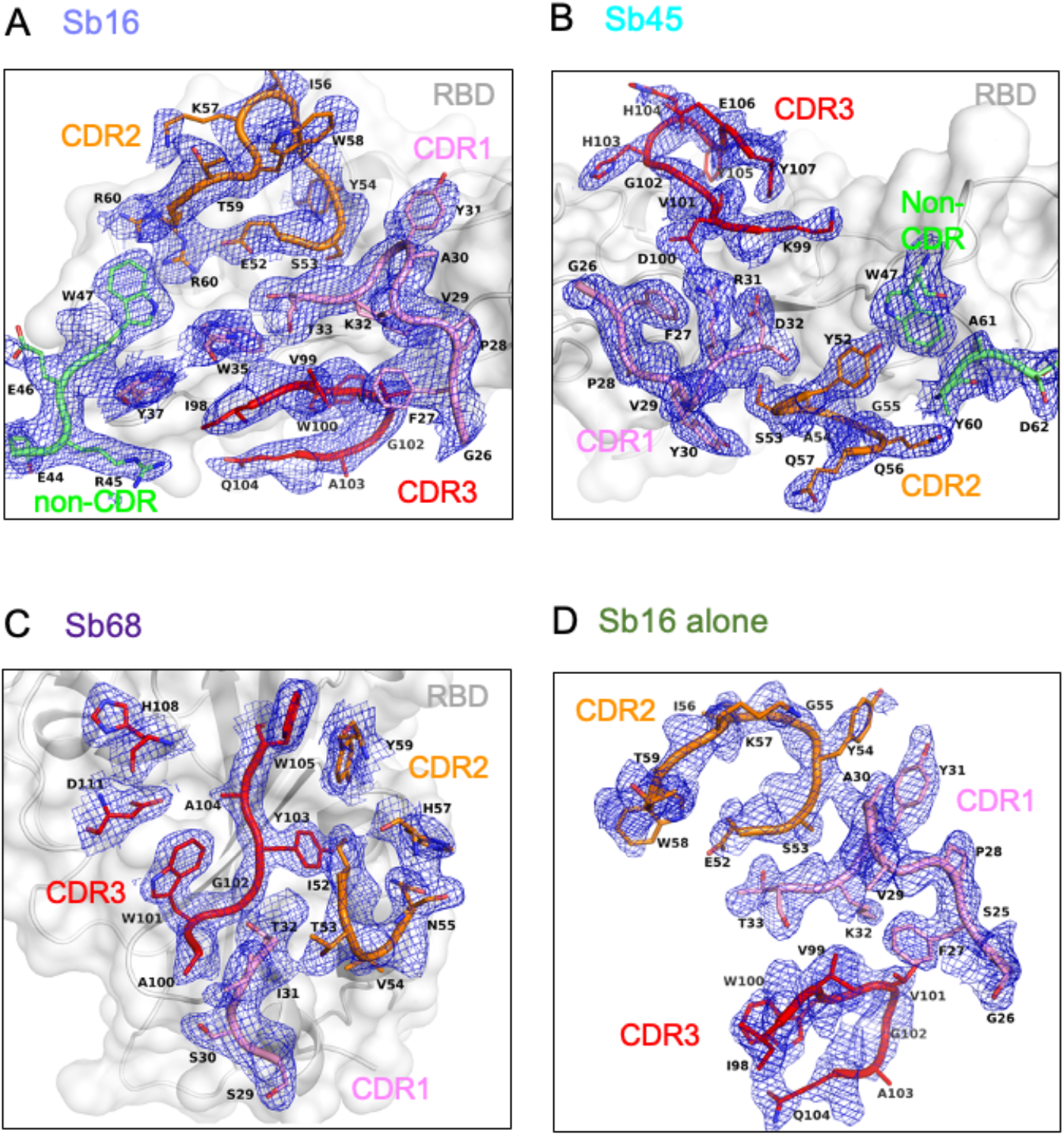
Electron density (as blue) maps (2mFo-DFc) for CDR loops and those residues in contact with RBD: (A) Sb16 on RBD surface, resolution=2.6 Å, *R*_free_=0.277; (B) Sb45 on RBD surface, resolution=2.3 Å, *R*_free_=0.216; (C) Sb68 on RBD surface, resolution=2.6 Å, *R*_free_=0.255; (D) Sb16 alone, resolution=2.1 Å, *R*_free_=0.259. Contour at 1.0σ, CDR1 loop as pink, CDR2 loop as orange, CDR3 loop as red, non-CDR residues as lime, and RBD is in background as gray.

**Fig. S5.**
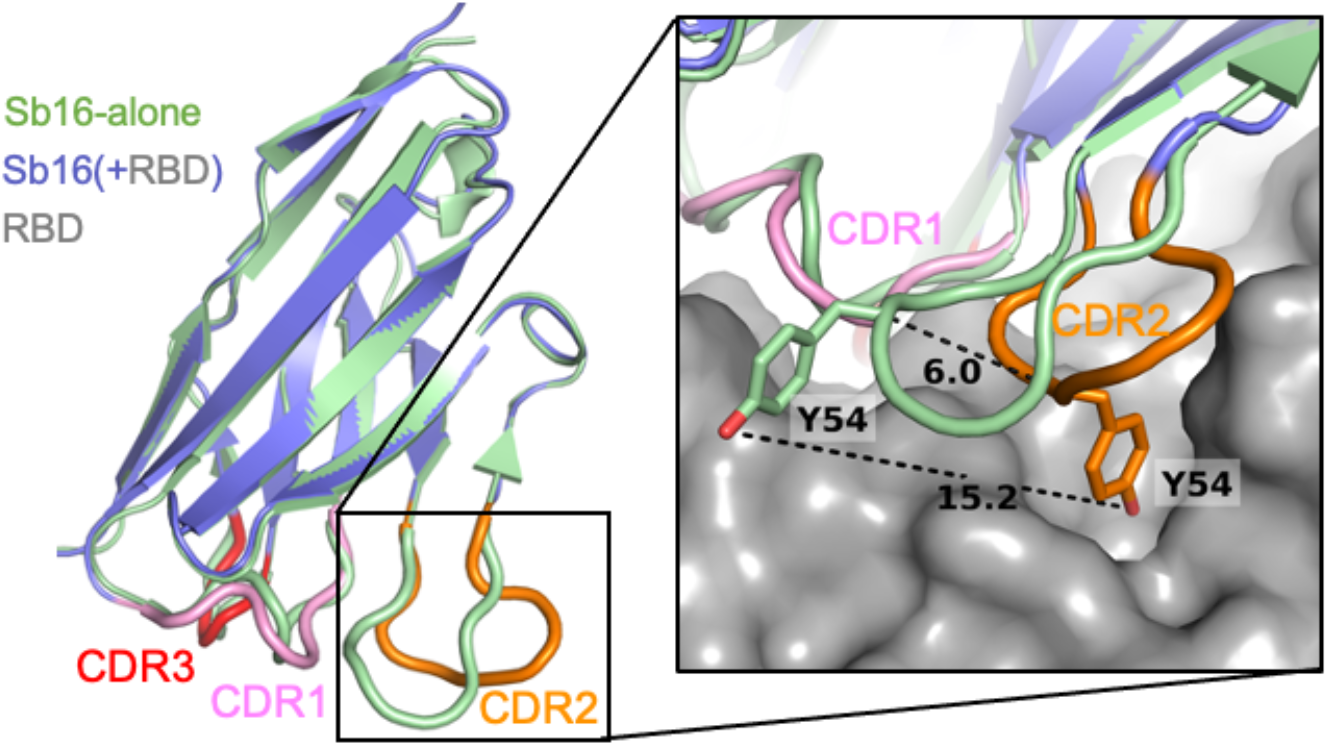
A superimposition of Sb16 alone (“unliganded”, free, green) and Sb16–RBD (“liganded”, complexed, slate) reveals the large movement of CDR2 loop (about 6 Å). Particularly, Y54 moved about 15 angstroms and dipped into a binding pocket which surrounded by epitopic residues of Q409, E406, D405, R403, G416, K417, I418, N422, L455, Y453, Y495. (RBD surface as gray).

**Fig. S6.**
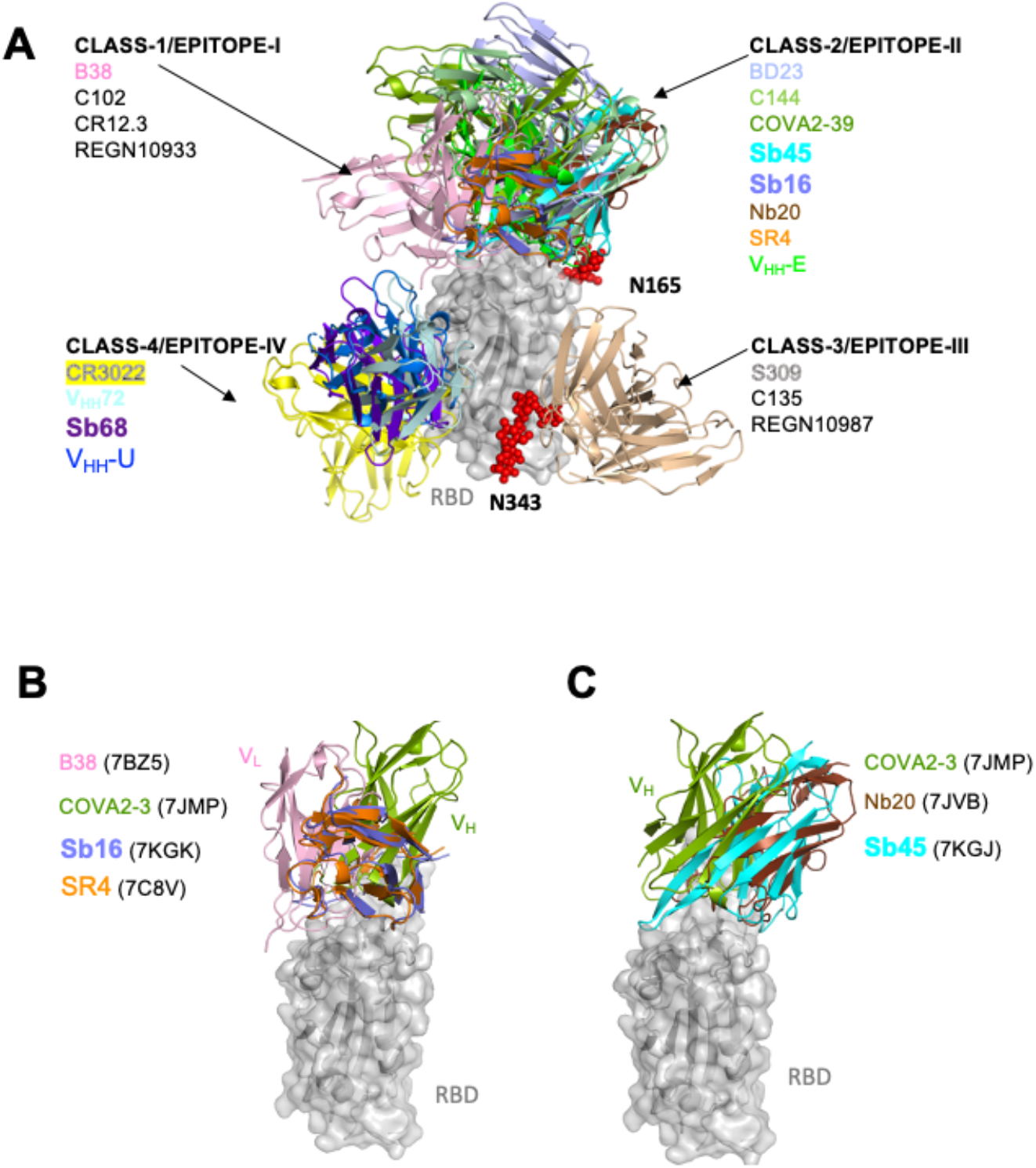
Sybodies, nanobodies, and antibodies bind different epitopic regions of the RBD. (A) Definition of Classes according to Barnes et al. (*31*) and Epitope area according to Xiang et al.(*12*). Epitope I to IV are almost the same as Class 1 to 4, except for Epitope V which spans Class 3 and Class 4 (not shown). Color codes are for representative Fab (only shown are the variable domains) or sybody/nanobody. Sb68 falls into Class-4/Epitope-IV overlapping with V_HH_72 and HH-E, and CR3022. RBD is in gray and two N-glycans (N165 and N343 in red) are shown. (B) Sb16 clashes with L-chain of B38 (7BZ5) of Class-1 and H-chain of COVA2-39 (7JMP) H-chain Class-2. It falls between Class-1/Epitope-I and Class-2/Epitope-II but almost covers SR4. (C) Sb45 overlaps with H-chain of COVA2-39 (7JMP) of Class-2 and lies in the same orientation as Nb20 (7JVB) - Epitope-II.

**Fig. S7.**
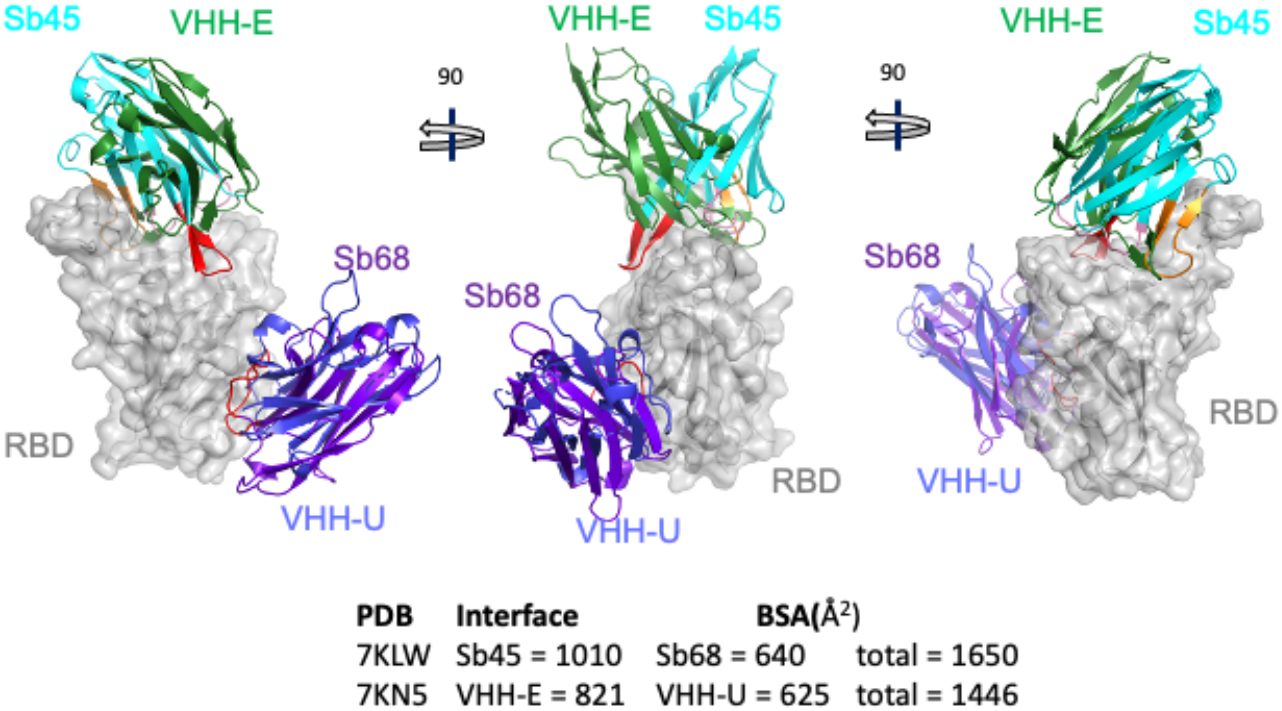
Comparison of the ternary sybody structure (Sb45–RBD–Sb68, 7KLW) with the ternary nanobody structure (V_HH_-E–RBD–V_HH_-U, 7KN5), Koenig et al. (*32*). Sb45 in cyan, Sb68 in purpleblue, RBD in gray, V_HH_-E in green, V_HH_-U in blue. CDR1, CDR2, and CDR3 of Sb45, CDR3 of Sb68 are highlighted in pink, orange, and red. Three views are each rotated by 90°. CDR3 and CDR2 loops of Sb45 ride along both sides of the RBD, while V_HH_-E uses only an extended CDR3 loop on the side. Sb68 is slightly lower than V_HH_-U while V_HH_-E is similar to V_HH_72.

**Fig. S8.**
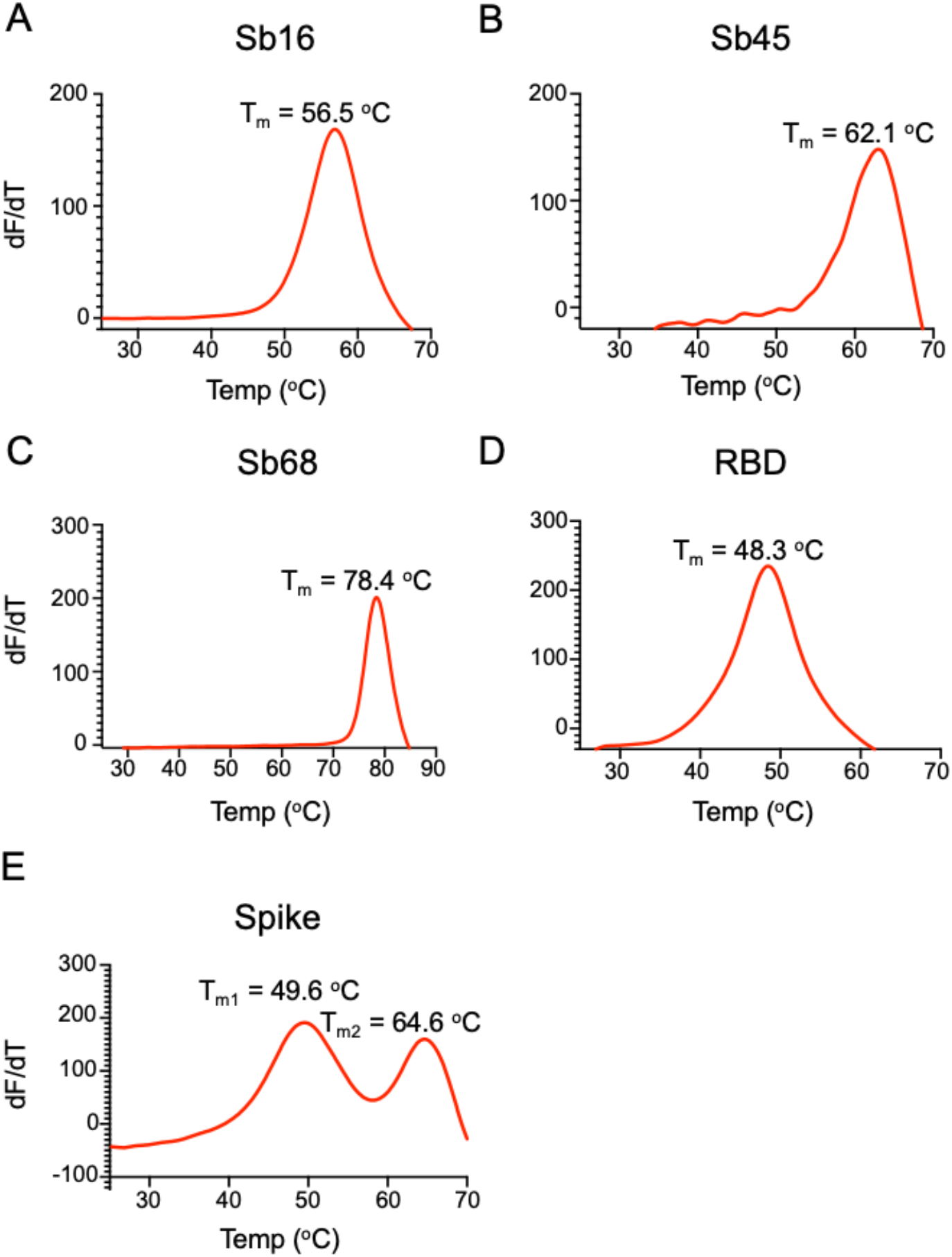
Sybodies, RBD, and spike protein reveal unique thermal stability. T_m_ of each of the indicated purified proteins was determined by thermal melt analysis as described in Materials and Methods. Note the biphasic behavior of the trimeric S protein.

**Table S1.**
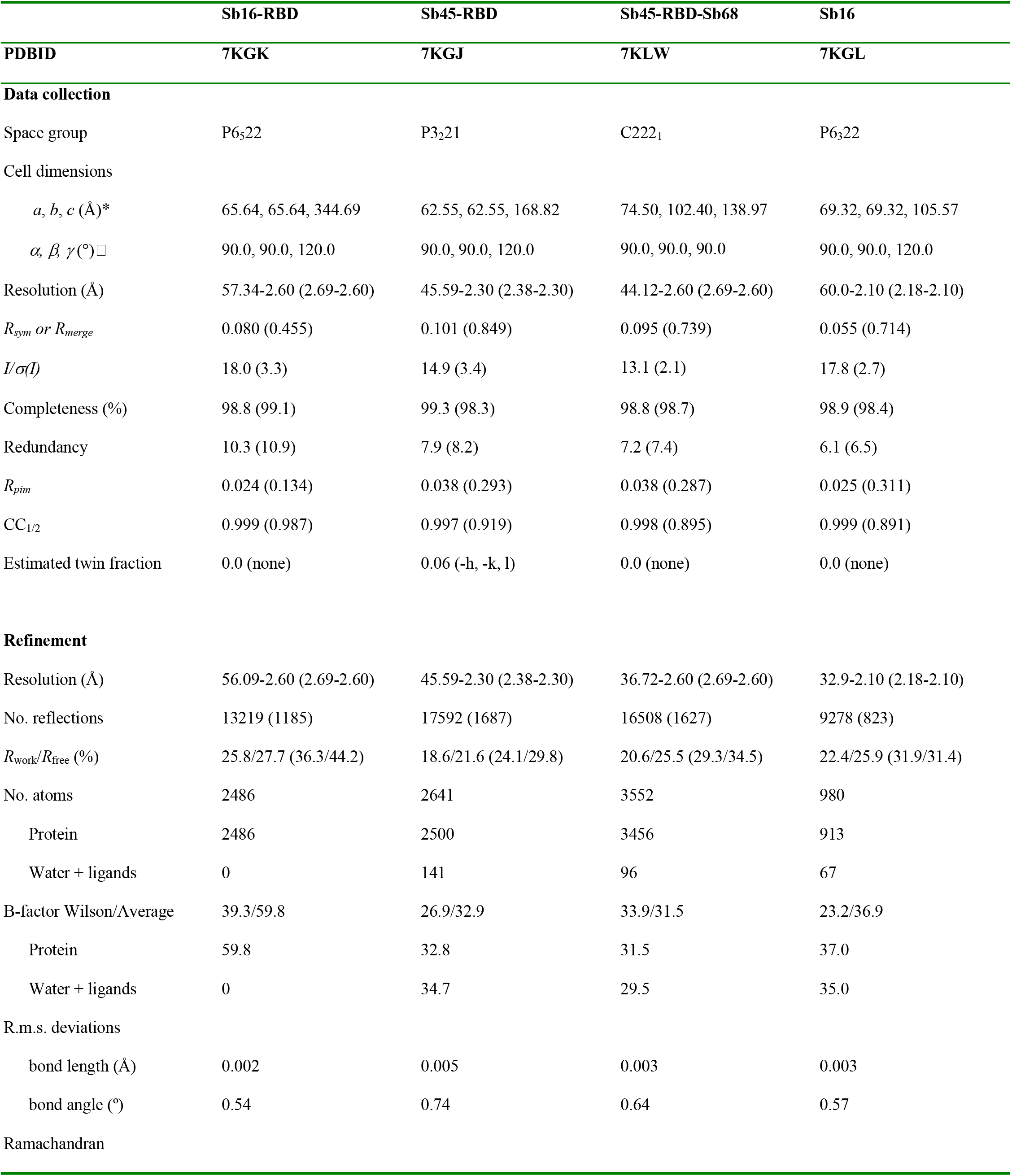
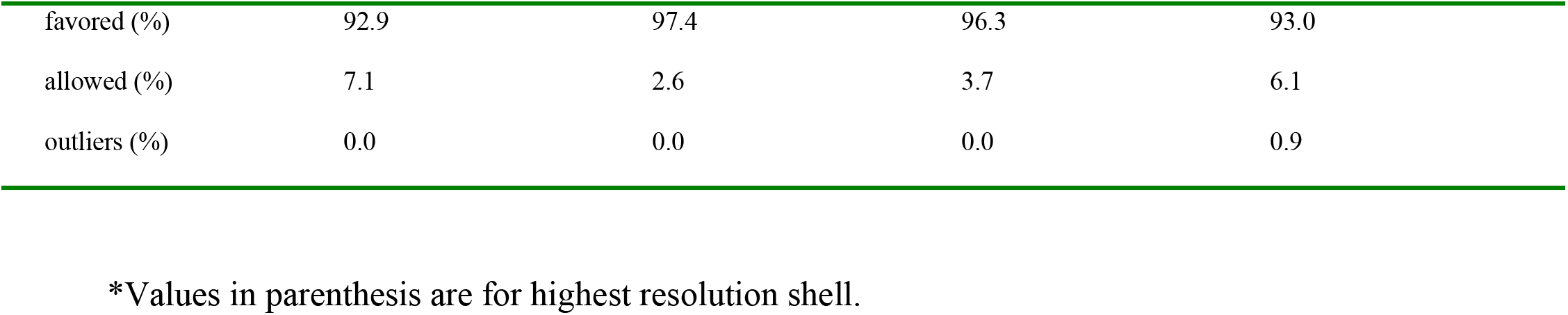
X-ray data collection and refinement statistics

**Table S2A.**
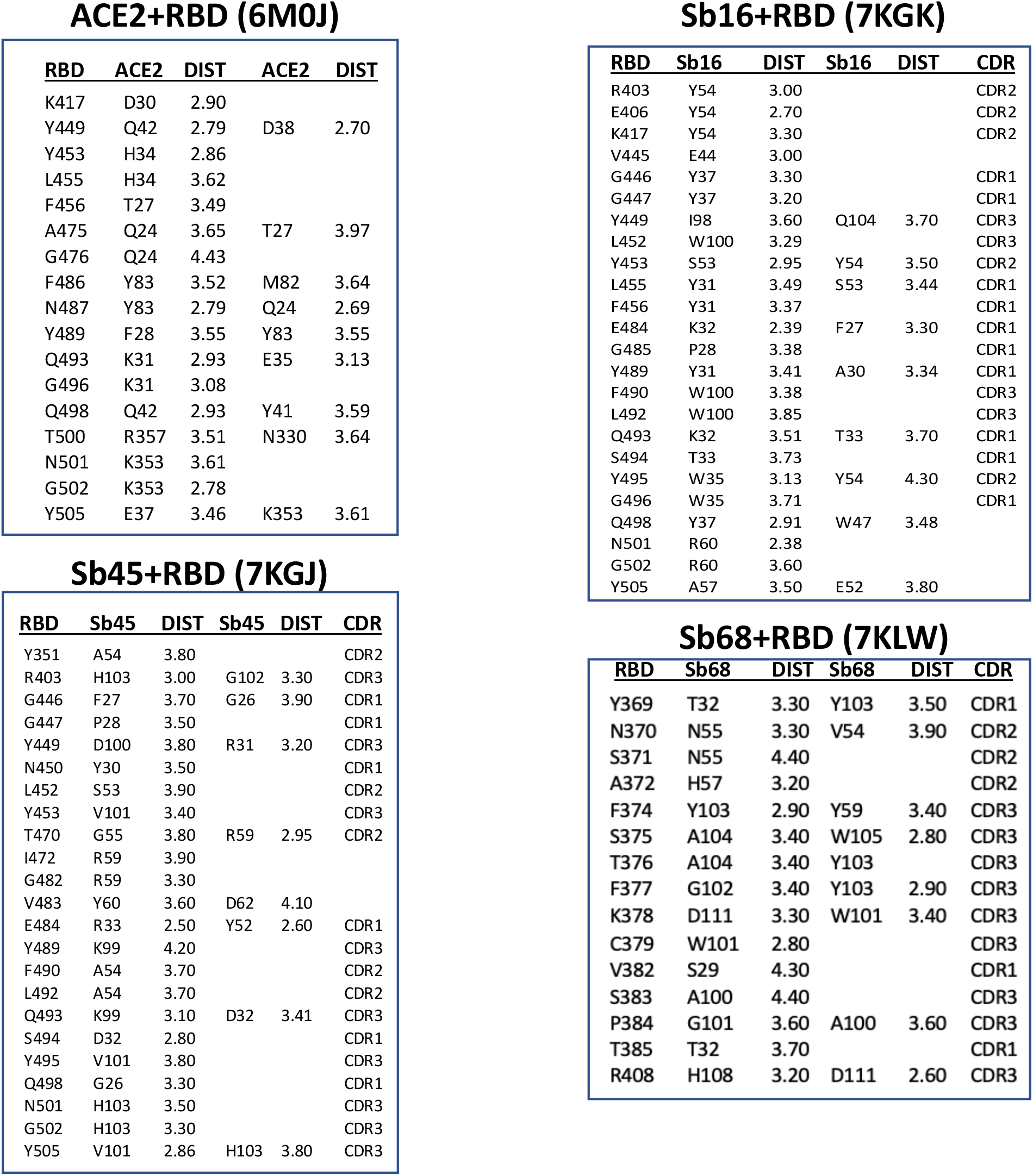
Interactions between RBD Residues and either ACE2 or sybody residues. In each rectangle (first column) is listed the identification of an RBD amino acid in contact with one or two residues of the indicted chain. Distance between residues is given in Å. For the sybodies, the particular CDR designation is also indicated.

ACE2+RBD (6M0J)
| RBD | ACE2 | DIST | ACE2 | DIST |
| --- | --- | --- | --- | --- |
| K417 | D30 | 2.90 |  |  |
| Y449 | Q42 | 2.79 | D38 | 2.70 |
| Y453 | H34 | 2.86 |  |  |
| L455 | H34 | 3.62 |  |  |
| F456 | T27 | 3.49 |  |  |
| A475 | Q24 | 3.65 | T27 | 3.97 |
| G476 | Q24 | 4.43 |  |  |
| F486 | Y83 | 3.52 | M82 | 3.64 |
| N487 | Y83 | 2.79 | Q24 | 2.69 |
| Y489 | F28 | 3.55 | Y83 | 3.55 |
| Q493 | K31 | 2.93 | E35 | 3.13 |
| G496 | K31 | 3.08 |  |  |
| Q498 | Q42 | 2.93 | Y41 | 3.59 |
| T500 | R357 | 3.51 | N330 | 3.64 |
| N501 | K353 | 3.61 |  |  |
| G502 | K353 | 2.78 |  |  |
| Y505 | E37 | 3.46 | K353 | 3.61 |

Sb16+RBD (7KGK)
| RBD | Sb16 | DIST | Sb16 | DIST | CDR |
| --- | --- | --- | --- | --- | --- |
| R403 | Y54 | 3.00 |  |  | CDR2 |
| E406 | Y54 | 2.70 |  |  | CDR2 |
| K417 | Y54 | 3.30 |  |  | CDR2 |
| V445 | E44 | 3.00 |  |  |  |
| G446 | Y37 | 3.30 |  |  | CDR1 |
| G447 | Y37 | 3.20 |  |  | CDR1 |
| Y449 | I98 | 3.60 | Q104 | 3.70 | CDR3 |
| L452 | W100 | 3.29 |  |  | CDR3 |
| Y453 | S53 | 2.95 | Y54 | 3.50 | CDR2 |
| L455 | Y31 | 3.49 | S53 | 3.44 | CDR1 |
| F456 | Y31 | 3.37 |  |  | CDR1 |
| E484 | K32 | 2.39 | F27 | 3.30 | CDR1 |
| G485 | P28 | 3.38 |  |  | CDR1 |
| Y489 | Y31 | 3.41 | A30 | 3.34 | CDR1 |
| F490 | W100 | 3.38 |  |  | CDR3 |
| L492 | W100 | 3.85 |  |  | CDR3 |
| Q493 | K32 | 3.51 | T33 | 3.70 | CDR1 |
| S494 | T33 | 3.73 |  |  | CDR1 |
| Y495 | W35 | 3.13 | Y54 | 4.30 | CDR2 |
| G496 | W35 | 3.71 |  |  | CDR1 |
| Q498 | Y37 | 2.91 | W47 | 3.48 |  |
| N501 | R60 | 2.38 |  |  |  |
| G502 | R60 | 3.60 |  |  |  |
| Y505 | A57 | 3.50 | E52 | 3.80 |  |

Sb45+RBD (7KGJ)
| RBD | Sb45 | DIST | Sb45 | DIST | CDR |
| --- | --- | --- | --- | --- | --- |
| Y351 | A54 | 3.80 |  |  | CDR2 |
| R403 | H103 | 3.00 | G102 | 3.30 | CDR3 |
| G446 | F27 | 3.70 | G26 | 3.90 | CDR1 |
| G447 | P28 | 3.50 |  |  | CDR1 |
| Y449 | D100 | 3.80 | R31 | 3.20 | CDR3 |
| N450 | Y30 | 3.50 |  |  | CDR1 |
| L452 | S53 | 3.90 |  |  | CDR2 |
| Y453 | V101 | 3.40 |  |  | CDR3 |
| T470 | G55 | 3.80 | R59 | 2.95 | CDR2 |
| I472 | R59 | 3.90 |  |  |  |
| G482 | R59 | 3.30 |  |  |  |
| V483 | Y60 | 3.60 | D62 | 4.10 |  |
| E484 | R33 | 2.50 | Y52 | 2.60 | CDR1 |
| Y489 | K99 | 4.20 |  |  | CDR3 |
| F490 | A54 | 3.70 |  |  | CDR2 |
| L492 | A54 | 3.70 |  |  | CDR2 |
| Q493 | K99 | 3.10 | D32 | 3.41 | CDR3 |
| S494 | D32 | 2.80 |  |  | CDR1 |
| Y495 | V101 | 3.80 |  |  | CDR3 |
| Q498 | G26 | 3.30 |  |  | CDR1 |
| N501 | H103 | 3.50 |  |  | CDR3 |
| G502 | H103 | 3.30 |  |  | CDR3 |
| Y505 | V101 | 2.86 | H103 | 3.80 | CDR3 |

**Table S2A.** Interactions between RBD Residues and either ACE2 or sybody residues.
| RBD | Sb68 | DIST | Sb68 | DIST | CDR |
| --- | --- | --- | --- | --- | --- |
| Y369 | T32 | 3.30 | Y103 | 3.50 | CDR1 |
| N370 | N55 | 3.30 | V54 | 3.90 | CDR2 |
| S371 | N55 | 4.40 |  |  | CDR2 |
| A372 | H57 | 3.20 |  |  | CDR2 |
| F374 | Y103 | 2.90 | Y59 | 3.40 | CDR3 |
| S375 | A104 | 3.40 | W105 | 2.80 | CDR3 |
| T376 | A104 | 3.40 | Y103 |  | CDR3 |
| F377 | G102 | 3.40 | Y103 | 2.90 | CDR3 |
| K378 | D111 | 3.30 | W101 | 3.40 | CDR3 |
| C379 | W101 | 2.80 |  |  | CDR3 |
| V382 | S29 | 4.30 |  |  | CDR1 |
| S383 | A100 | 4.40 |  |  | CDR3 |
| P384 | G101 | 3.60 | A100 | 3.60 | CDR3 |
| T385 | T32 | 3.70 |  |  | CDR1 |
| R408 | H108 | 3.20 | D111 | 2.60 | CDR3 |

**Table S2b.**
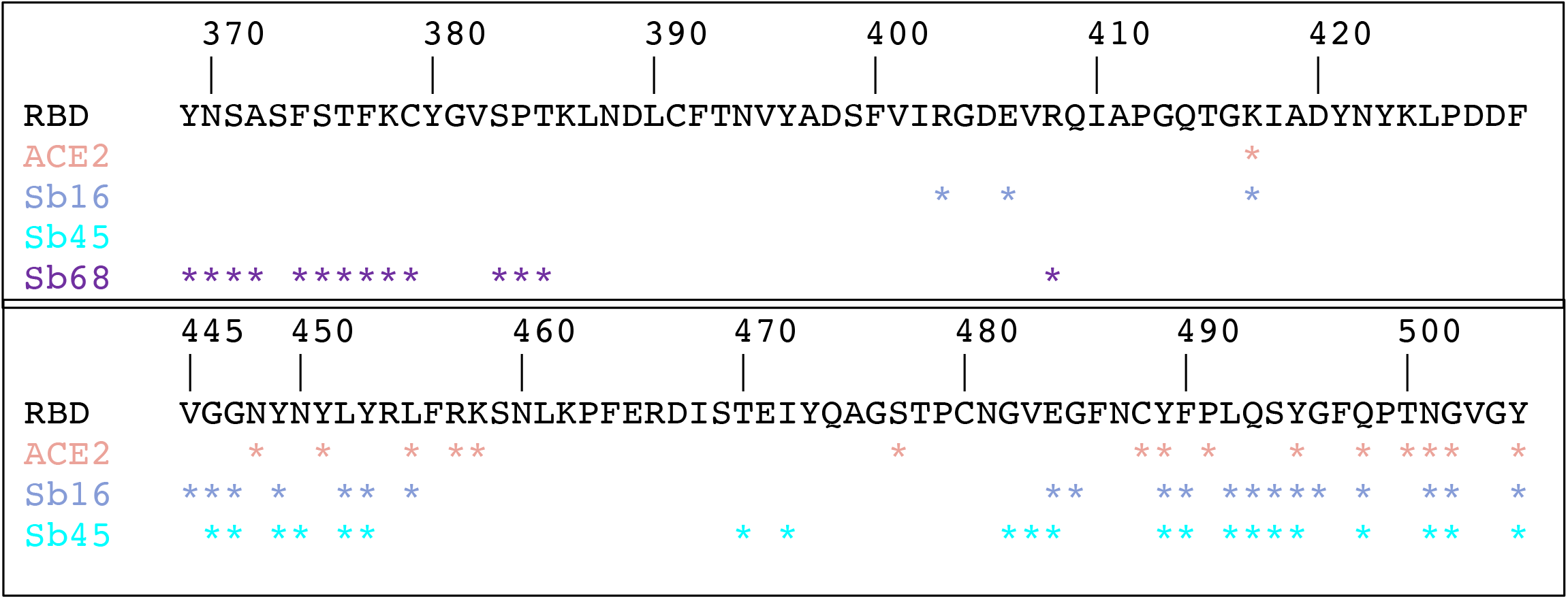
Tabulation of residues of RBD that contact each of the indicated chains, based on 6M0J for ACE2–RBD, 7KGK for Sb16-RBD, 7KGJ for Sb45-RBD, and 7KLW for the Sb68-RBD interface. Note that ACE2 contacts broadly overlap those for Sb16 and Sb45, but that Sb68 contacts are distinct.

**Table S3.** Variety of Buried Surface Area (BSA) at interfaces of Sybodies, Nanobodies, and Fabs with RBD or Spike. BSA of each of indicated interface was calculated from the relevant chains using PISA (https://www.ebi.ac.uk/pdbe/pisa/), to account for the non-overlapping surface area. BSA in Å^2^. N_res_ indicates the number of residues in contact; N_HB_, the number of potential hydrogen bonds; N_SB_, the number of potential salt bridges.

| **Complex (A + B)** | **BSA($\text{\AA}^2$)** | **$N_{\text{res}}^{\text{A}}$** | **$N_{\text{res}}^{\text{B}}$** | **$N_{\text{HB}}$** | **$N_{\text{SB}}$** | **PDB** | **Res. $\text{\AA}$** |
| --- | --- | --- | --- | --- | --- | --- | --- |
| **ACE2+RBD** | **844** | **26** | **26** | **13** | **2** | **6M0J** | 2.45 |
| Sb16+RBD | **1003** | 26 | 29 | 8 | 1 | 7KGK | 2.60 |
| Sb45+RBD | **976** | 27 | 33 | 15 | 4 | 7KGJ | 2.10 |
| *(Sb45)+RBD | **1010** | 26 | 35 | 14 | 4 | 7KLW_B | 2.60 |
| *(Sb68)+RBD | **640** | 21 | 17 | 9 | 4 | 7KLW_C | 2.60 |
| *(Sb45+Sb68)+RBD | **1650** | 47 | 52 | 23 | 8 | 7KLW | 2.60 |
| *(VHH-E)+RBD | **821** | 29 | 27 | 13 | 1 | 7KN5_C | 1.87 |
| *(VHH-U)+RBD | **625** | 16 | 20 | 17 | 0 | 7KN5_E | 1.87 |
| *(VHH-E+VHH-U)+RBD | **1446** | 45 | 47 | 30 | 1 | 7KN5 | 1.87 |
| H11D4+RBD | **599** | 20 | 20 | 11 | 4 | 6YZ5 | 1.80 |
| H11H4+RBD | **637** | 17 | 19 | 5 | 4 | 6ZBP | 1.85 |
| (CR3022)+RBD | **991** | 19 | 20 | 7 | 4 | 6XC7_HL | 2.88 |
| VHH72+RBD | **796** | 21 | 25 | 9 | 2 | 6WAQ_A | 2.20 |
| Nb20+RBD | **705** | 22 | 21 | 9 | 4 | 7JVB | 3.29 |
| Nb6+Spike | **788** | 24 | 21 | 9 | 1 | 7KKK_D | 3.03 |
| Sb23+Spike | **772** | 21 | 22 | 5 | 0 | 7A25_D | 3.06 |
| Sb23+Spike | **585** | 18 | 21 | 5 | 0 | 7A29_D | 2.94 |
| Ty1+Spike | **795** | 23 | 26 | 3 | 0 | 6ZXN_D | 2.93 |
| C144+Spike | **689** | 22 | 24 | 7 | 0 | 7K90_H | 3.24 |
| C002+Spike | **728** | 22 | 23 | 7 | 1 | 7K8S_H | 3.40 |
| *(REGN10933)+RBD | **935** | 28 | 30 | 9 | 1 | 6XDG_BD | 3.90 |
| *(REGN10987)+RBD | **607** | 21 | 19 | 5 | 0 | 6XDG_AC | 3.90 |
| *(REGN10933+ | 1542 | 49 | 49 | 5 | 0 | 6XDG | 3.90 |
| REGN10987)+RBD |  |  |  |  |  |  |  |

